# Using machine learning to predict protein-protein interactions between a zombie ant fungus and its carpenter ant host

**DOI:** 10.1101/2022.09.09.507359

**Authors:** Ian Will, William C. Beckerson, Charissa de Bekker

**Affiliations:** University of Central Florida, Department of Biology, 4110 Libra Drive, 32816, Orlando, FL, USA; Utrecht University, Department of Biology, Microbiology, Padualaan 8, 3584 CH, Utrecht, NL

## Abstract

Parasitic fungi produce proteins that modulate virulence, alter host physiology, and trigger host responses. These proteins, classified as a type of “effector,” often act via protein-protein interactions (PPIs). The fungal parasite *Ophiocordyceps camponoti-floridani* (zombie ant fungus) manipulates *Camponotus floridanus* (carpenter ant) behavior to promote transmission. The most striking aspect of this behavioral change is a summit disease phenotype where infected hosts ascend and attach to an elevated position. Plausibly, interspecific PPIs drive aspects of *Ophiocordyceps* infection and host manipulation. Machine learning PPI predictions offer high-throughput methods to produce mechanistic hypotheses on how this behavioral manipulation occurs. Using D-SCRIPT to predict host-parasite PPIs, we found ca. 6,000 interactions involving 2,083 host proteins and 129 parasite proteins, which are encoded by genes upregulated during manipulated behavior. We identified multiple overrepresentations of functional annotations among these proteins. The strongest signals in the host highlighted neuromodulatory G-protein coupled receptors and oxidation-reduction processes. We also detected *Camponotus* structural and gene-regulatory proteins. In the parasite, we found enrichment of *Ophiocordyceps* proteases and frequent involvement of novel small secreted proteins with unknown functions. From these results, we provide new hypotheses on potential parasite effectors and host targets underlying zombie ant behavioral manipulation.

## Introduction

Fungal parasites employ a diverse array of secreted molecules to defend themselves, promote infection, and modify their hosts. In certain cases, infection can even lead to parasitic manipulation of host behavior. Often termed “effectors,” secreted fungal molecules play a critical role in host-parasite dynamics that include both widely shared and highly specific mechanisms ^1–4^. Parasite effectors have been suggested to play key roles during the infection of insects by entomopathogenic fungi through their interactions with host nucleic acids, carbohydrates, lipids, small metabolites, and proteins ^3, 5–7^. Bioinformatic exploration of effector protein biology offers a high-throughput method to develop mechanistic hypotheses of how some parasites can modify the behavior of their hosts. Here, we use such approaches to investigate the behavior-manipulating fungus *Ophiocordyceps camponoti-floridani* (Florida zombie ant fungus) and its insect host *Camponotus floridanus* (Florida carpenter ant).

Myrmecophilous *Ophiocordyceps* fungi are typically species-specific parasites that have co-evolved with their ant hosts over millions of years ^8^. This close relationship has resulted in fungus-ant interactions that alter host behaviors in ways that are adaptive for the parasite. Manipulated host ants succumb to a summit disease, affixing themselves to elevated positions and dying at locations that promote fungal growth and transmission ^9–13^. Behavioral changes preceding this final summit could include host-adaptive responses, parasite-adaptive manipulations, or general symptoms of disease that span hyperactivity, uncoordinated foraging, decreased nestmate communication, and convulsions ^9, 14–17^. While these modified ant behaviors have been observed in nature for some time ^18^, explorations of the fungal effectors involved are relatively more recent endeavors ^14, 19–25^. These multiple “-omics” studies have provided hypotheses about the parasite and host molecules that play a role in establishing the behavioral phenotypes observed, but leave their potential interactions open to interpretation.

A large number of fungal molecules and ant pathways may be involved in *Ophiocordyceps* behavioral manipulation, including neuro-modulators and-protectants, insect hormones, feeding, locomotion, circadian rhythms and light-sensing, and muscular hyperactivity^14, 17, 19, 21, 22, 25–29^. Fungal proteins such as bacterial-like enterotoxins, protein tyrosine phosphatases, peptidases (e.g., S8 subtilisin-like serine proteases), and undescribed small secreted proteins (uSSPs) have been proposed as mechanisms of manipulation that presumably function by PPIs. These hypotheses are supported by strong gene upregulation of these candidates during active manipulation of the ant and genomic comparisons among the *Ophiocordyceps* ^14, 19, 24^. Numerous other such proteins are supported as plausible effectors as well. Testing them *in vivo* will be a costly and laborious process in an emerging, non-traditional model organism such as *O. camponoti-floridani*. New evidence linking previously hypothesized candidate proteins and pathways to host-parasite protein-protein interactions (PPIs) would give strong support to select top predictions for functional validation.

Leveraging big-data and computational methods to explore host-parasite biology is a continuing and multifaceted effort. Yet, examples of using these assets in PPI prediction remain limited ^30, 31^. Proteome-scale bioinformatic prediction of PPIs has largely been employed to describe protein interaction networks within a single organism, as was done for the host species, *C. floridanus* ^32^. However, PPI predictions in some interspecific host-parasite relationships have been made, including animal or plant hosts and bacterial, viral, or fungal parasites ^33–36^. Predicting PPIs is typically done using either an interolog or machine learning method. Interologs are orthologous interactions conserved across species; each protein in one PPI has orthologs in the second PPI ^37^. While a useful prediction tool, detecting an interolog in a new system requires established reference PPIs from other systems. This constraint by available data would result in a very limited number of predictions when analyzing interactions between non-model organisms. Machine-learning methods offer a more flexible alternative. In particular, neural network deep learning tools for PPI prediction have become increasingly available in recent years and continue to be an active field of development ^38–41^. Here, predicted PPIs are still informed by a known set of PPIs used for training data. However, once trained, the program is able to predict novel interactions outside the original dataset ^35, 42^. The deep learning tool D-SCRIPT has shown an uncommon flexibility in cross-species PPI predictions when trained on one organism but tested on a different species ^35^. Protein embeddings that encode putative structural information are used to train D-SCRIPT models, which have produced predicted protein-protein contact maps that largely agree with experimentally measured docked proteins ^35, 36, 43^. Although other machine learning methods can outperform D-SCRIPT in same-species contexts, the cross-species generalizability of D-SCRIPT’s predictive power is an important step for research on non-model or multi-species systems ^35, 38^. Notably, *Ophiocordyceps* and other fungi secrete a range of taxonomically distinct uSSPs that are often thought to be effectors in host-parasite interactions ^14, 19, 24, 44–46^. Therefore, bioinformatic techniques that hinge upon well described protein annotations or are subject to strong species-specific biases can only have limited success in assigning uSSPs to PPIs.

For our analyses between *O. camponoti-floridani* and *C. floridanus* proteins, we anchor our reporting on group-level functional enrichments of predicted positive PPIs. We chose this approach in consideration of the reported D-SCRIPT default model error rates on insect (*Drosophila*) and fungal (*Saccharomyces*) species. In these tests, predicted positive PPIs contained 71-79% true-positives (precision rate) and captured 22-36% of all true-positives (recall rate) ^35^, which indicate this tool is strongest in broad characterizations rather than validating specific, previously hypothesized PPIs. Similarly, we benchmarked D-SCRIPT on experimentally supported fungus-animal interspecific PPIs and found comparable low true-positive recall coupled with higher precision. The authors of D-SCRIPT have also suggested that this tool can outperform an earlier deep learning method in reproducing gene ontology (GO) term enrichments of experimentally identified virus-human PPIs ^35^.

We used D-SCRIPT to predict possible *O. camponoti-floridani* and *C. floridanus* PPIs involved in behavioral manipulation. We filtered these predictions into subsets we considered most likely to include relevant interactions. As such, our study focuses on PPIs involving putatively secreted *O. camponoti-floridani* proteins encoded by genes that were previously found to be upregulated during manipulated summiting ^19^. These fungal proteins are plausibly exported to the host environment during manipulation, providing support for hypotheses implicating such proteins in manipulation of host behavior.

We highlight aspects of functional enrichments after filtering out *Camponotus* protein interactions with fungi that do not naturally infect or manipulate this host (“aspecific” fungi). These aspecific fungi cover different lifestyles and phylogenetic relationships to *Ophiocordyceps*: (*i*) *Cordyceps bassiana* (i.e., *Beauveria bassiana*), a generalist entomopathogen in the order Hypocreales (which includes *Ophiocordyceps*), (*ii*) *Trichoderma reesei*, a plant-degrading saprophyte in the order Hypocreales (although the genus also includes entomopathogens), and (*iii*) *Saccharomyces cerevisiae*, a phylogenetically distant saprophytic yeast in the order Saccharomycetales with a smaller secretome. This approach brought the unique *Ophiocordyceps* PPIs to the forefront but does not discount the aspecific predictions. Rather, it brings additional attention to a narrower set of specific PPIs that could include co-evolved adaptations underlying *Ophiocordyceps*-manipulated *Camponotus* behavior.

## Materials and Methods

### Fungus-animal PPI benchmarking

The Host-Pathogen Interaction Database (v 3.0) provided experimentally supported PPIs between 5 fungal and 12 animal species. We used this data to benchmark D-SCRIPT in predicting interspecific PPIs. The types of experimental support we selected included direct interactions and reactions (e.g., “phosphorylation reaction”) but excluded associations, colocalizations, and genetic effects (e.g., “additive genetic interaction”). We only included PPIs with proteins ≤ 2,000 amino acids, for computational reasons. We further limited similar PPIs to include only one representative interaction by clustering all proteins in the dataset by 40% similarity using MMseqs2 (v. 9-d36de) (single step easy-clustering with minimum sequence identity 0.4) ^47^. For any PPIs that connected the same two homologous protein clusters, we selected one of those PPIs at random and removed the others. In total, this produced 567 experimentally verified fungus-animal PPIs.

In addition to these verified, positive fungus-animal PPIs, we constructed a set of negative interactions to use in evaluating D-SCRIPT. Previously, D-SCRIPT evaluation datasets have used randomly paired proteins from the tested organism as negative PPI examples ^35^. In this vein, we created artificial interspecific fungus-animal PPIs using random yeast (*S. cerevisiae*) and fly (*Drosophila melanogaster*) proteins collected from the STRING database (v. 11.0) ^48^. Rather than randomly pairing proteins from all organisms in the fungus-animal dataset, we chose these two organisms because they both: (*i*) were represented in the positive PPIs, (*ii*) had robust PPI data available for removing known positive interactions from the random pairings, (*iii*) were used in the original D-SCRIPT evaluations, and (*iv*) match *Ophiocordyceps* and *Camponotus* as a fungus-insect pair ^35^. Using the aforementioned 40% clustering method, we filtered out any random yeast-fly PPIs that were similar to positive fungus-animal interactions. We also removed any that were similar to known, high-confidence, experimentally-supported fly or yeast PPIs listed in the STRING database (“experiments score” > 0 and “combined score” ≥ 700) ^48^. Then, we kept only one representative of PPIs that were similar to each other. After filtering, we selected 5,670 random PPIs (10-fold the number of positive cases) to use as negative examples in the evaluation dataset.

We used D-SCRIPT (v 0.1.5) with the default human-protein pretrained model and settings in evaluation mode with this fungus-animal dataset (567 positive and 5,670 negative PPIs) ^35^. D-SCRIPT assigns each tested protein interaction an edge value (i.e., a confidence score to rank possible PPIs) with a default threshold for positive prediction of ≥ 0.5, on a scale of zero to one. These parameters have been previously used to predict PPIs within fly and yeast ^35^.

### Protein sequences and initial selection

To predict PPIs between *O. camponoti-floridani* secreted proteins and *C. floridanus* proteins, we retrieved sequence information for both organisms from high-quality genome assemblies integrating long-read technology (GenBank accessions GCA_012980515.1 and GCA_003227725.1, respectively) ^19, 49^. For *Camponotus* proteins, we used the sequence represented by the single longest isoform (n = 12,512). We predicted *Ophiocordyceps* proteins to have a localization outside of the cell by combining eight annotation tools (SignalP v6.0, TMHMM, Phobius, Prosite, PredGPI, NucPred, and TargetP v2.0) ^50–56^ following the relatively conservative approaches outlined in Beckerson et al., 2019 ^1^. We focused on secreted proteins (n = 586) and excluded transmembrane proteins (n = 1,301) in our downstream analyses, as we considered these to most plausibly constitute effectors. Moreover, our preliminary findings showed that transmembrane proteins were generally less specific in their binding (Supplementary Discussion S1). All remaining non-secreted, non-transmembrane proteins were considered intracellular (n = 5,568). We annotated secreted proteins as uSSPs if they were under 300 amino acids in length and lacked any BLAST description, GO term, or PFAM domain (n = 154) ^19^. We used proteins ≤ 2,000 amino acids due to computational constraints. Limiting the length of input proteins removed 63 (1.1%) intracellular *Ophiocordyceps* proteins and 299 *Camponotus* proteins (2.4%).

### PPI predictions

We used D-SCRIPT (v 0.1.5) in prediction mode with the default human-protein pretrained model and settings to predict PPIs (as above for fungus-animal benchmarking) ^35^. We tested every PPI combination between secreted *O. camponoti-floridani* proteins and the entire *C. floridanus* proteome (Fig. 1 step 1).

**Figure 1.**
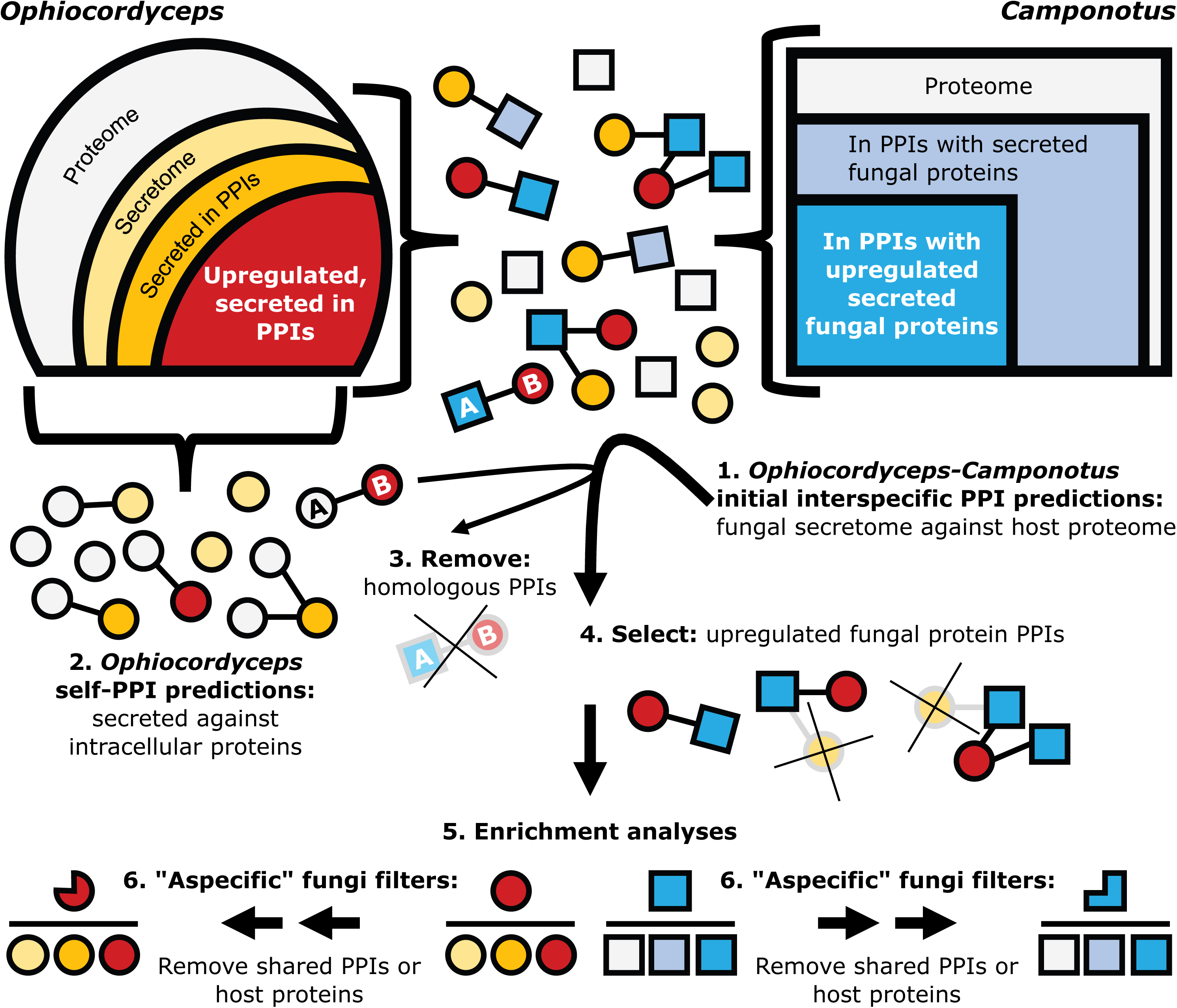
Conceptual framework for PPI testing, selection, and analysis. Step 1) We tested every secreted *O. camponoti-floridani* protein (secretome) with every *C. floridanus* host protein (proteome). Steps 2, 3) If an interspecific PPI and an *Ophiocordyceps* self-interaction PPI shared a common fungal protein paired with a protein homologous between both species, we removed that PPI from our predictions. Step 4) Subsequently, we filtered predicted PPIs to focus on those that involved fungal proteins encoded by genes that were upregulated during manipulation. Step 5) We analyzed host and parasite proteins from these PPIs with hypergeometric enrichment analyses. For enrichment analyses, the background protein set was either the parasite secretome (*Ophiocordyceps*) or host proteome (*Camponotus*). Step 6) We also performed enrichment analyses after removing PPIs or host proteins that were predicted in interactions between aspecific fungi and *C. floridanus*. Fungal proteins are shown as circles and ant proteins as squares. Proteins are color-coded, with *Ophiocordyceps* upregulated, secreted proteins in interspecific PPIs indicated in red and their interacting *Camponotus* proteins in bright blue. Shades of orange represent other secreted fungal protein categories while blue-gray indicates other host proteins in PPIs. All other proteins are shown in gray. Proteins “A” are *Ophiocordyceps* and *Camponotus* homologs, and protein “B” is a single fungal protein.

To identify self-interacting *Ophiocordyceps* PPIs, we tested secreted *Ophiocordyceps* proteins against intracellular *Ophiocordyceps* proteins (Fig. 1 step 2). We then identified reciprocal homologs (putative orthologs) between *Ophiocordyceps* and *Camponotus* proteins using Proteinortho (v 5.0) with default settings ^57^. We filtered out PPIs that contained a fungal secreted protein predicted to bind both one of *Ophiocordyceps*’ own intracellular proteins and a *Camponotus* protein that was orthologous to that fungal intracellular protein (Fig. 1 step 3). We reasoned that an interspecific PPI would more likely indicate an interaction related to infection or manipulation when the *Ophiocordyceps* protein was predicted to only bind a *Camponotus* protein, without also binding a fungal ortholog to that ant protein. We considered this to be a favorable tradeoff with possibly overlooking parasite effectors that may target well-conserved biological processes.

We narrowed the remaining PPIs to those involving secreted fungal proteins encoded by differentially expressed genes (DEGs) (Fig. 1 step 4). These genes were upregulated during manipulated summiting behavior in *C. floridanus* ants experimentally infected with *O. camponoti-floridani*, as compared to their expression in culture ^19^. By incorporating empirical gene expression data, we anchored our computational analyses in predictions that are most biologically relevant during manipulation. Although the chance exists that some parasite effectors act during manipulation without increased gene transcription, we presumed that in most cases, gene expression would be informative.

### Hypergeometric enrichment analyses

We performed hypergeometric enrichment analyses on proteins from PPIs with secreted, upregulated *Ophiocordyceps* proteins that only had predicted interactions with the host. For enrichment analyses of the fungal proteins in these PPIs, we used the *O. camponoti-floridani* secretome as the background. For enrichment analyses of the ant proteins in these PPIs, we used the *C. floridanus* proteome as the background (Fig. 1, step 5).

We examined annotated GO terms (biological processes and molecular functions combined) ^58^, PFAM domains ^59^, and weighted gene co-expression network analysis (WGCNA) ^60^ module memberships reported in the transcriptomics study from which we also obtained the above-mentioned DEGs ^19^. These WGCNA modules correlated expression of gene-networks to sample types in that study, i.e., (*i*) healthy ant controls, (*ii*) *in vitro* fungus controls, (*iii*) active fungus-manipulated ants, and (*iv*) dying fungus-manipulated ants. These modules were additionally correlated directly between *Ophiocordyceps* gene modules and *Camponotus* gene modules ^19^. We referenced their previously assigned names following the convention of ant-module-1 (A1) and fungus-module-1 (F1) as used in that study ^19^. For *Camponotus* proteins, we also tested for enrichments of DEG class (i.e., upregulated or downregulated gene expression from healthy controls to active manipulated ants) ^19^. For *Ophiocordyceps* proteins, we additionally tested for enrichment of uSSP annotations.

We performed the hypergeometric enrichment analyses with the R package timecourseRNAseq (v 0.0.9000) using default settings (significant enrichment at FDR ≤ 0.05) in R studio (v 2021.09.2) with R (v 4.1) ^61–63^. We plotted GO terms in semantic space to cluster related terms based on R code generated from REVIGO but did not remove any GO terms (R package ggplot2, v 3.3.5) ^64, 65^.

### Aspecific fungus-*Camponotus* PPI combinations

For more selective analyses of species-specific *Ophiocordyceps*-*Camponotus* PPIs, we produced aspecific PPIs from the secretomes of other fungi tested against the *C. floridanus* proteome. Removal of more common PPIs, leaving only those potentially unique to *Ophiocordyceps*, provided a view that was easier to interpret in the context of specialized parasite-host interactions, such as behavioral manipulation. However, a PPI need not be unique to a certain interspecific interaction to be relevant ^1^. Additionally, computationally predicted protein binding does not indicate that all such interactions occur in dynamic physiological environments. In and around the cell, protein localization, transcriptional control, post-translational modification, competing PPIs, and biochemical conditions (e.g., pH) could affect the likelihood of a predicted PPI to occur. The cellular environments within fungal cells and insect cells could plausibly differ in many ways, shaping protein activity. Therefore, we used the removal of aspecific interactions to help us focus on some of the, likely, more important PPIs involved in establishing *Ophiocordyceps* manipulation of ant behavior, but still anchored our analyses and discussion on the larger dataset.

We chose three fungi based on lifestyle, phylogeny, and availability of well-annotated genomes to generate aspecific PPIs: *C. bassiana* (GenBank accession GCA_000280675.1), *T. reesei* (GenBank accession GCA_000167675.2), and *S. cerevisiae* (GenBank accession GCF_000146045.2) ^7, 66, 67^. As for *Ophiocordyceps* (see above), we predicted each fungal secretome and removed secreted proteins over 2,000 amino acids long, which only removed one *T. reesei* protein (0.002%). The secretomes of *C. bassiana*, *T. reesei*, and *S. cerevisiae* fungi included 833, 505, and 153 proteins, respectively. These proteins were then paired with the *Camponotus* proteome for D-SCRIPT predictions (see above).

To find orthologous proteins between *Ophiocordyceps* and each of the aspecific fungi we used Proteinortho (v 6.0) with default settings while enforcing blastp+ and single-best reciprocal hits (i.e., -sim=1) ^57^. Using this information, we identified shared PPIs between *Ophiocordyceps* and *Camponotus*, and aspecific fungi and *Camponotus*; i.e., interactions between orthologous fungal proteins and the same ant protein (Supplementary Fig. S1). More stringently, we also applied a filter that eliminated shared host proteins with aspecific fungal interactions, regardless of orthology. This resulted in a set of *Ophiocordyceps* PPIs where the *Camponotus* protein was never predicted in any aspecific PPI (Supplementary Fig. S1). After removing either shared PPIs or shared host proteins, we again performed enrichment analyses (see above) to investigate overrepresented annotations among the *Ophiocordyceps*-specific PPIs (Fig. 1 step 6). To visualize changes in PPI connectivity and abundance that resulted from these aspecific filters, we used Cytoscape (v 3.8.2) to produce networks of host proteins and their respective *Ophiocordyceps* binding partners ^68^.

## Results

### Fungus-animal benchmarking PPIs

To evaluate D-SCRIPT’s performance on predicting interspecific PPIs between fungi and animals, we used experimentally supported interactions from the Host-Pathogen Interaction Database ^69^. Our evaluation dataset included 567 of these verified, positive interactions from a diversity of fungal and animal species. We combined these with a 10-fold negative background (5,670 PPIs) constructed by randomly pairing fungal (*S. cerevisiae*) and insect (*D. melanogaster*) proteins. For this evaluation with a 1 positive:10 negative ratio test data, the model performance metric, area under precision-recall curve (AUPR), has a baseline of 0.091. D-SCRIPT produces results well above this level for fly-only ^35^, yeast-only ^35^, and these fungus-animal data (AUPR 0.552, 0.405, and 0.373, respectively) (Table 1). Although interspecific interactions indeed appear to be challenging to predict, our benchmarking data (Table 1) suggest D-SCRIPT still provides meaningful insights in interspecific contexts.

**Table 1.** Fungus-animal PPI D-SCRIPT benchmark. We evaluated D-SCRIPT on interspecific fungus-animal PPIs using positive interactions from the Host-Pathogen Interaction Database (n = 567) and negative interactions from randomly paired yeast and fly proteins (n = 5,670). For this dataset and those published with D-SCRIPT ^35^ with 1 postive:10 negatives, the baseline AUPR is 0.091. Recall is the proportion of true positive examples in the dataset that the model correctly predicted to interact. Precision is the proportion of predicted positives that were indeed correct. Based on these benchmarking data, interspecific PPIs are more difficult for D-SCRIPT to predict, but, we suggest, they still can be detected at rates useful for initial biological investigation and characterization of group-level trends.

| Evaluation dataset | AUPR | Precision | Recall |
| --- | --- | --- | --- |
| Fly, as published <sup>35</sup> | 0.552 | 0.798 | 0.359 |
| Yeast, as published <sup>35</sup> | 0.405 | 0.706 | 0.223 |
| Fungus-animal | 0.373 | 0.720 | 0.169 |

### Ophiocordyceps-Camponotus PPIs

We tested a total of 7,156,818 potential interspecific PPIs between the parasite secretome and host proteome (Fig. 1 step 1). Of these possible protein-protein combinations, 0.33% were positively predicted as PPIs (n = 23,629 PPIs) (Table 2, Supplementary Fig. S2). This result is in line with a previous 0.95% positive D-SCRIPT prediction rate from a single-species dataset testing 50 million potential PPIs ^35^.

**Table 2.**
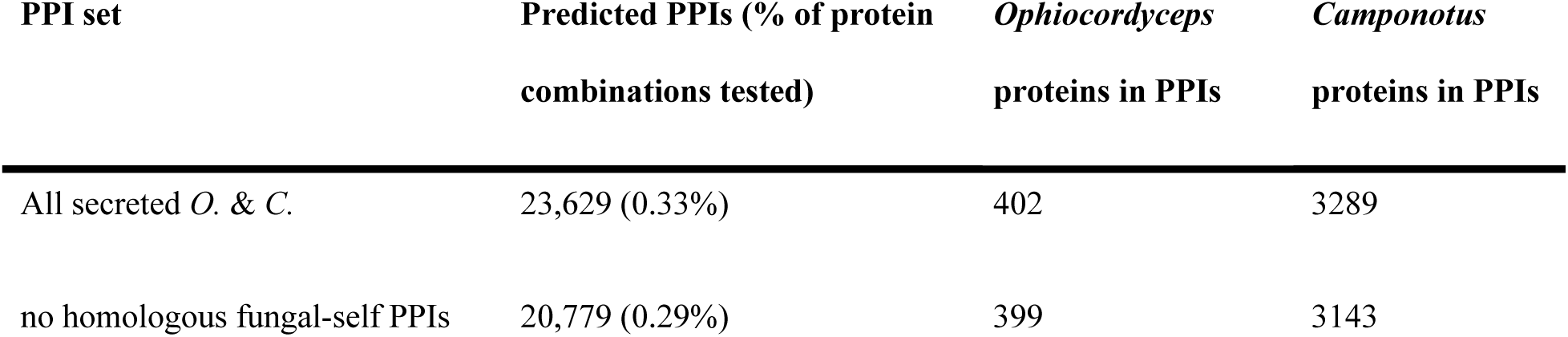
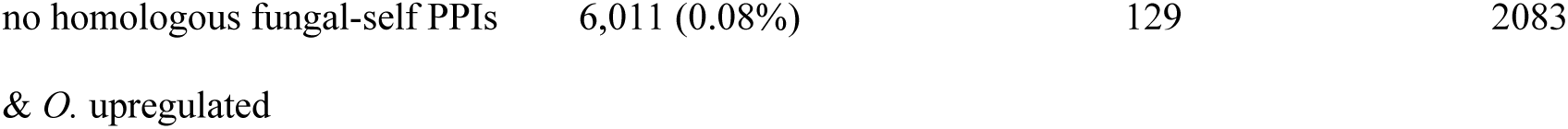
PPI sets overview. We tested 7,156,818 combinations of secreted *O. camponoti-floridani* (*O.*) proteins (n = 586) with every *C. floridanus* (*C.*) protein (n = 12,213). Predicted PPIs were combinations returned with an edge value of ≥ 0.5. We then created PPI subsets based on homologous interaction between secreted and intracellular fungal proteins and fungal gene upregulation during manipulation.

From these positively predicted interspecific interactions, we removed 2,850 PPIs that were homologous to within-*Ophiocordyceps* interactions (Fig. 1 step 2 and 3, Table 2). We then selected those involving *Ophiocordyceps* proteins encoded by upregulated DEGs during manipulation. This resulted in 6,011 predicted PPIs. We used this set of putative effector-host PPIs, containing 129 unique *Ophiocordyceps* proteins and 2,083 unique *Camponotus* proteins, for further analyses (Fig. 1 step 4, Table 2, Supplementary File S1).

### PPIs between aspecific fungi and *Camponotus* in comparison to *Ophiocordyceps*

The proportions of positive PPI predictions between aspecific fungi and *C. floridanus* (range 0.26-0.28%) and secreted proteins predicted to be in at least one PPI (range 68-76%) were similar to the values observed for *Ophiocordyceps* (0.33% and 69%, respectively) (Table 2, Supplementary Table S1). Eliminating orthologous PPIs between *Ophiocordyceps* and any of the aspecific fungi (i.e., shared PPIs) removed 1,776 of the 6,011 (30%) *Ophiocordyceps*-*Camponotus* PPIs (Table 3, Supplementary Fig. S1). Eliminating PPIs involving host proteins that were also predicted to be bound by aspecific fungi, regardless of orthology of the fungal binding partner (i.e., shared host proteins), removed 1,936 of 2,083 (93%) host proteins, leaving only 157 *Ophiocordyceps*-*Camponotus* PPIs (3%) (Table 3, Supplementary Fig. S1). In line with secretome size, phylogeny, and entomopathogenic lifestyle, *C. bassiana* shared the most overlap with *Ophiocordyceps* predictions (Supplementary Fig. S1). As expected, removing PPIs involving shared host proteins led to major changes in enrichment results while removing shared PPIs led to more modest changes (Table 3, Supplementary File S2). We primarily report the results based on all three fungi that we included in this study. However, those based on *C. bassiana* alone were largely similar (Supplementary Fig. S1, Supplementary File S2). While these filters assisted us in ordering and emphasizing results (i.e., signals maintained through the strictest filters are being prioritized), we did not use this approach to wholly select PPIs of interest. Results presented below only use aspecific filters where stated.

**Table 3.** Annotation enrichments of PPI proteins. We performed enrichment analyses on interspecific PPIs without homology to *Ophiocordyceps* self-interactions and involving upregulated, secreted *Ophiocordyceps* proteins and any *Camponotus* proteins (Fig. 1 step 5) with or without additional filtering based on shared aspecific fungal interactions (Fig. 1 step 6). We found no enrichment signal for uSSPs in the fungus. Only in the unfiltered analysis were *Camponotus* DEGs overrepresented – for both upregulated and downregulated genes. The number of gene co-expression modules enriched are given under column WGCNA. While typically more stringent filtering reduced the number of enrichments detected, it increased the number of enriched GO terms for *Ophiocordyceps*. These enrichments included oxidation-reduction terms and various general binding terms (e.g., “binding,” “catalytic activity,” “cofactor binding,” or “ion binding”).

| Test proteins (n) | GO | PFAM | WGCNA |
| --- | --- | --- | --- |
| *Ophiocordyceps* (129) | 0 | 1 | 3 |
| after shared PPIs removed (125) | 1 | 1 | 3 |
| after shared host proteins removed (26) | 11 | 0 | 2 |
| *Camponotus* (2,083) | 50 | 74 | 9 |
| after shared PPIs removed (1,901) | 44 | 64 | 9 |
| after shared host proteins removed (147) | 12 | 3 | 2 |

### *Ophiocordyceps* enrichment signals

We found that *Ophiocordyceps* proteins in PPIs with *Camponotus* proteins were enriched for one PFAM domain and three WGCNA modules (Fig. 2, Table 3).

**Figure 2.**
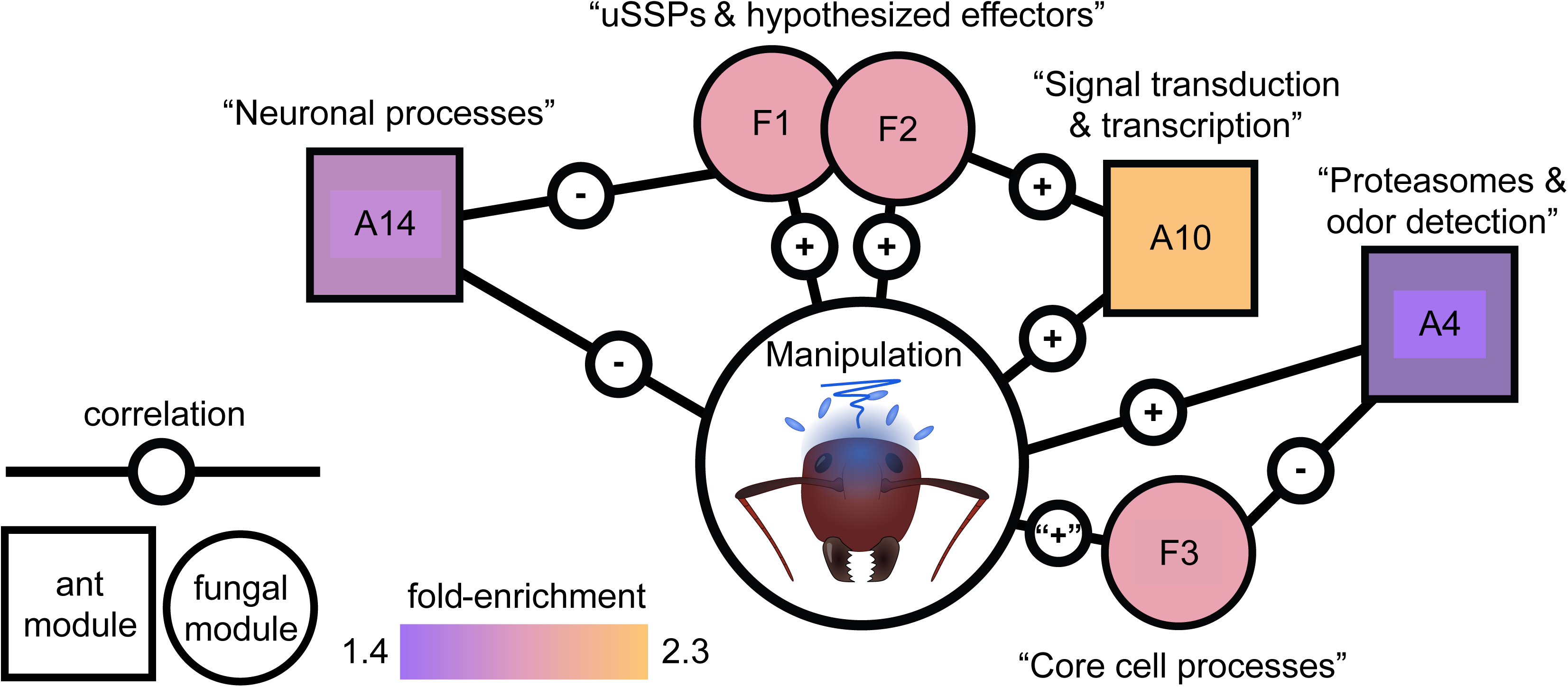
Enriched WGCNA modules and their correlations to manipulation and each other. Module membership, correlations, and functional summary of modules were based on previous work ^19^. Each module is a mutually exclusive network of genes that are coexpressed across control, manipulated ant, and dying ant conditions. Here, we only depict modules enriched among PPIs and only their correlations to the manipulated state or between host-parasite modules directly. Core cell processes module F3 was clearly negatively correlated to control conditions, with modest positive correlations divided between both live manipulation and dying ants ^19^, as indicated by “+” here. While module F1 and F3 were also notably correlated to neuronal process module A15, this ant module is not depicted here as it was not enriched among the PPIs detected in this study.

### S8 serine peptidases

The enriched PFAM domain was a “peptidase S8 domain,” indicating an overrepresentation of fungal serine proteases that could cleave host proteins. The five fungal peptidase S8 proteins in this enrichment were predicted to interact with 34 host proteins across 39 PPIs (Supplementary File S1). Among these host proteins, there were seven kinesin-like proteins (motor proteins), five nuclear pore proteins, and a (pro-)resilin that is an insect cuticle and connective tissue protein ^70^. Even after removing PPIs that were shared with aspecific fungi, the enrichment signal for S8 peptidases was retained. However, the much more stringent removal of shared host targets eliminated the S8 peptidase enrichment result (Fig. 1 step 6, Table 3).

### Fungal manipulation WGCNA modules

We found fungal WGCNA modules F1, F2, and F3 to be enriched (Fig. 2). We previously reported modules F1 and F2 to have significant positive correlations to manipulation. This means that most of the fungal genes in those modules were upregulated, in a similar co-expressed manner, during manipulated summiting behavior. As such, these modules contained multiple genes hypothesized to mediate infection or manipulation processes (“uSSPs and hypothesized effectors” modules) (Fig. 2). With fungal gene expression clearly negatively correlated with control cultures, module F3 was modestly positively correlated to fungi in both manipulated hosts and dying hosts. This module largely contained genes related to “core cell processes” in the fungus (e.g., DNA packaging) (Fig. 2). All three fungal modules also had significant direct correlations to ant WGCNA modules, with F1 being negatively correlated to ant modules A14 and A15 – both associated with host neuronal processes (Fig. 2). Core cell processes module F3 was positively correlated with neuronal process module A15 ^19^.

The overrepresentations of all three fungal WGCNA modules was retained after eliminating shared PPIs, but uSSP and effectors module F1 was lost after removing shared host proteins (Table 3).

### *Camponotus* enrichment signals

We found that *Camponotus* PPI proteins were enriched for 50 GO terms, 74 PFAM domains, and nine WGCNA modules (Table 3). As GO term and PFAM domain enrichments often indicated similar functions, we present results largely from the GO term perspective. We highlight PFAM domains for emphasis or to discuss biologically interesting results not well captured by GO terms alone.

### G protein-coupled receptor signaling

We detected GO term enrichments for “G protein-coupled receptor (GPCR) activity” and “GPCR signaling pathway” (Fig. 3). These enrichments resulted from 71 GPCR-related PPIs with 16 unique *Ophiocordyceps* proteins (Supplementary File S1). All but two receptor proteins in these PPIs carried the PFAM domain “7 transmembrane receptor (rhodopsin family).” We found 33 unique GPCR-related protein genes (of 128 possible with these GO terms), with a range of putative ligands: 11 neuropeptides (n = 12 receptors), four biogenic monoamines (dopamine, serotonin, octopamine, tyramine) (n = 11 receptors), acetylcholine (n = 2 receptors), a spider venom toxin protein (alpha-latrotoxin) (n = 1 receptor), and opsin blue-and ultraviolet-sensitive receptors (n = 2 receptors). Fourteen of these 33 *Camponotus* receptors (or receptor subunits) belonged in ant-neuronal processes WGCNA modules A14 and A15, which showed overall reduced expression during manipulation compared to healthy ants (i.e., the modules were negatively correlated to manipulation) ^19^. Host GPCR-related PPI proteins were predicted to bind 16 fungal proteins that included five uSSPs, a carboxylesterase, a glycosyl hydrolase, a CAP cysteine-rich secretory protein, and a protein tyrosine phosphatase. Most of these fungal proteins were previously assigned to fungal uSSPs and effectors module F2 (n = 9) but included some in modules F1-4 (Supplementary File S1), which all were also correlated to infection and manipulation of hosts.

**Figure 3.**
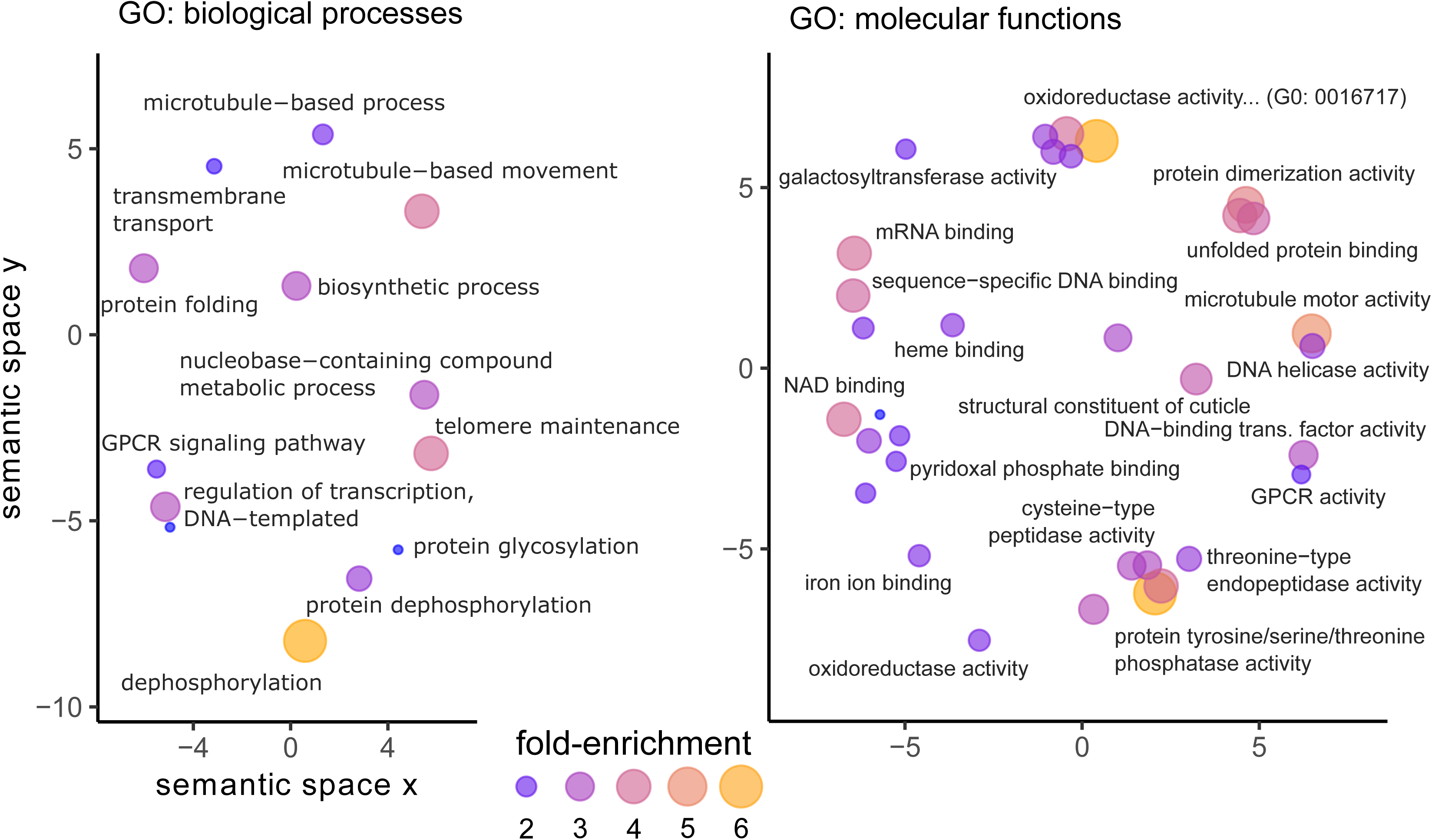
Enrichments for *C. floridanus* proteins in PPIs with upregulated secreted fungal proteins. GO terms are plotted in semantic space, which lacks any inherent unit or value other than to cluster terms by functional similarity. Some labels were omitted for readability and when not relevant to our discussion of the results.

The enrichment signals for GPCRs were retained after we removed both PPIs and host proteins shared with aspecific fungi (Fig. 4). Twenty-eight of the 33 GPCR host proteins remained after removing shared PPIs. However, only seven remained after filtering out shared host proteins. These host proteins, predicted to be uniquely bound by *Ophiocordyceps* during manipulation, were a putative dopamine receptor 2, Oamb octopamine receptor, methuselah-like 10 receptor, orexin receptor, trissin receptor, cAMP-dependent protein kinase subunit, and receptor without an informative BLAST description (“uncharacterized protein”). The four *Ophiocordyceps* proteins binding these host receptors were predicted in many PPIs (range = 101 to 458). The dopamine receptor 2, Oamb octopamine receptor, and uncharacterized receptor shared the same fungal uSSP partner. The orexin and trissin receptors also shared an unannotated secreted fungal protein (larger than an uSSP).

**Figure 4.**
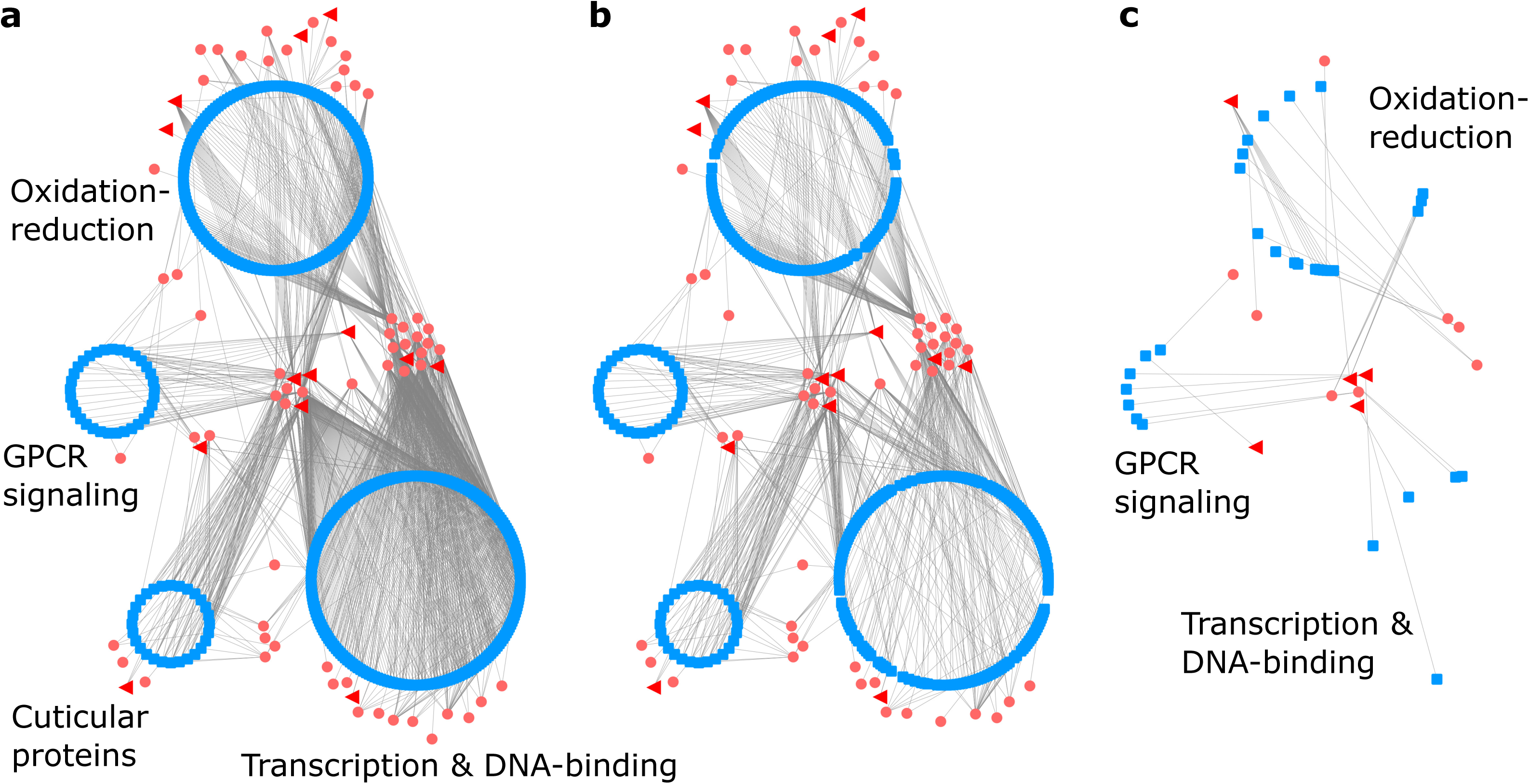
PPI network with host proteins contributing to selected GO term enrichments. Host proteins (blue square nodes) are clustered and labeled by their general GO term functional category and connected (gray line edges) to their fungal PPI partners that are either secreted proteins (small pink circle nodes) or uSSPs (large red triangle nodes) that were found to be upregulated during *Ophiocordyceps* manipulation of *Camponotus* behavior. Proteins may have PPI connections outside of those depicted here. a) PPI network without aspecific PPI filtering. b) Network of remaining PPIs after removing shared PPIs with aspecific fungi. Although nearly 30% of PPIs (edges) were removed, most proteins (nodes) and enrichment results were kept. c) Network of remaining PPIs after removing PPIs with all host proteins that also interacted with aspecific fungi, regardless of orthology of the fungal binding partner. GO terms related to G-protein coupled receptor signaling and some oxidation-reduction enrichments persisted after this filter. Enrichment signals for transcription and DNA-binding were lost, but some individual proteins remained.

### Oxidation-reduction

We also found the enrichment of oxidation-reduction functions among *Camponotus* proteins in PPIs (six supporting GO terms) (Fig. 3, Supplementary File S1). Altogether, 159 host proteins had oxidation-reduction annotations, forming 321 PPIs with 48 parasite proteins. A plurality of the host proteins were encoded by cytochrome P450 genes (70 unique genes, 15 putative subtypes). This abundance of cytochrome P450, a hemoprotein, also contributed to the enrichment of “heme-binding” and “iron ion binding” GO terms (Fig. 3) and the PFAM domain “cytochrome P450” (Supplementary File S1). These *Camponotus* cytochrome P450 proteins paired with 10 unique *Ophiocordyceps* genes, consisting of uSSPs and fungal cytochrome P450s. Other notable contributors to the host enrichment of oxidation reduction related functions were peroxiredoxins, laccases, dehydrogenases, and a vat-1 synaptic vesicle membrane-like protein (Supplementary File S1). The 48 fungal proteins included cytochrome P450s, oxidases and oxidoreductases, amidases, peptidases, and an enterotoxin.

Enrichment signals for four of the six oxidation-reduction GO terms were retained after we removed PPIs that were shared with the aspecific fungi. Only two remained after we applied the more stringent shared host protein filter (Supplementary File S2). Additionally, indicators of cytochromes, heme and iron binding GO terms and the “cytochrome P450” PFAM domain, continued to be enriched after either filter (Supplementary File S2). Most host proteins passed the shared PPI filter (140 of 159), but only 19 remained after removing shared host proteins (Fig. 4). Cytochrome P450 proteins shifted from a plurality to a dominant majority after shared host protein filtering (17 of 19 proteins), with eight cytochrome P450 subtypes: 3A19, 315a1, 4C1, 49a1, 6a13, 6a14, 6k1, and 9e2.

### Transcription and DNA-binding

Multiple enrichments that can be associated to gene transcription and regulation were enriched. We found 204 host proteins annotated for any of “DNA binding,” “DNA-binding transcription factor activity,” “sequence-specific DNA binding,” “DNA helicase activity,” “regulation of transcription, DNA-templated,” and “mRNA binding” GO terms (Fig. 3). These proteins similarly contributed to the high enrichment of PFAM domains such as “helix-loop-helix DNA-binding domain,” “zinc Finger C4 type,” “‘paired box’ domain,” and “bZIP transcription factor” (Supplementary File S1). These 204 host proteins interacted with 43 fungal proteins to generate 922 PPIs. Among the 204 host proteins, we found transcription factors and regulators with many functions, which included examples associated with insect behavior and neuronal function such as Mushroom body large type Kenyon specific protein 1 (e.g., elevated in worker bees, foraging behavior), Gooseberry (e.g., neuromuscular junction function), Hairy (e.g., juvenile hormone mediated gene repression), photoreceptor specific nuclear receptor (light sensing), CLOCK (circadian clocks), and steroid and ecdysone nuclear receptors and induced proteins ^71–75^. Other proteins, most clearly understood from the neuronal development perspective thus far, produce atypical behavioral or locomotor phenotypes when dysregulated and include Jim Lovell, Dead ringer, Hairy/enhancer-of-split related, Achaete-scute complex proteins, and Forkhead ^76–81^. These 34 proteins taken as biologically relevant examples account for 186 of the 922 gene regulation PPIs, interacting with 20 *Ophiocordyceps* proteins covering a similar breadth of protein types as the 43 parasite proteins overall (Supplementary File S1). The 43 fungal proteins included five uSSPs that were also found in the PPIs reported above (Fig. 4A), three that were not, and various hydrolases, oxidoreductases, peptidases, and carboxylesterases.

All of the aforementioned GO terms and PFAM domains continued to be enriched after removing shared PPIs (191 of 204 host proteins retained), but none of these annotations continued to be enriched after removing shared aspecific host proteins (five host proteins retained) (Fig. 4, Supplementary File S2).

### Cuticular proteins

We found 28 ant proteins that shared the enriched GO term “structural constituent of cuticle” (Fig. 3) and PFAM domain “insect cuticle protein” (Supplementary File S1). We predicted these ant proteins interacted with 22 fungal proteins to form 138 PPIs (Fig. 4A). These host proteins included (pro-)resilin, endocuticle structural proteins, and a range of “cuticle proteins” (e.g., “cuticle protein 7”). The fungal proteins included a CAP protein and a carboxylesterase also predicted to bind GPCRs (see above). Four fungal uSSPs were shared with the transcription and DNA-binding host proteins. We additionally identified a glycosyl hydrolase, a disintegrin/reprolysin-like peptidase, a S8 subtilisin-like serine peptidase, M43 peptidases, an uSSP, and an unannotated secreted protein (larger than an uSSP). These enrichments were only kept under the aspecific shared PPI filter (with all 28 original cuticular proteins) and lost with the shared host protein filter (none of the 28 proteins were uniquely bound by *Ophiocordyceps*) (Fig. 4, Supplementary File S2).

### Ant DEGs and WGCNA modules

*Camponotus* genes that were previously determined to be differentially expressed during manipulation, as well as host gene modules correlated to manipulation, were enriched among PPI proteins (Supplementary File S1). Both up-and downregulated host DEGs were enriched. The upregulated host DEGs were enriched for gene/DNA regulatory functions and the downregulated DEGs for oxidation-reduction (Supplementary File S1). Three ant WGCNA modules were enriched that had significant correlations to manipulation, with an additional six that did not (of 22 possible modules total). “Neuronal processes” module A14 was negatively correlated to manipulation and fungal uSSP and effectors module F1 (Fig. 2). This host module included many genes putatively associated with neuronal function, including GPCRs reported above ^19^. The other host neuronal processes module, A15, was not enriched for PPIs. Module A10 was positively correlated to manipulation and contained enrichment signals for GO terms indicating “signal transduction and transcription” ^19^ (Fig. 2). Module A4 had a positive correlation to manipulation and contained enrichment signals for “proteasomes and odor detection” ^19^ (Fig. 2). All three of these host modules passed the shared PPI filter but did not persist after removing shared host targets with aspecific fungi.

## Discussion

The interspecific molecular interactions during manipulation of *C. floridanus* by *O. camponoti-floridani* are likely mediated by parasite effector proteins that target host proteins. To predict such PPIs, we used the machine learning tool D-SCRIPT, which infers structural relationships between proteins without relying on protein-wide sequence homology to known interactions. This flexibility allows D-SCRIPT models to extend predictions across organisms and beyond previously documented PPIs. We benchmarked D-SCRIPT against a dataset including experimentally supported fungus-animal PPIs. This evaluation indicated D-SCRIPT can produce meaningful predictions on interspecific PPIs, although it performs better on single-species data. D-SCRIPT predicted several thousand PPIs between *Camponotus* host proteins and upregulated, secreted *Ophiocordyceps* parasite proteins. Even with low recall (identification of true PPIs) functional enrichments and trends at the group level can plausibly be detected. Correct PPIs predictions reflect a true biological signal while, most likely, incorrect-prediction noise would not frequently produce coherent enrichments signals.

Due to the unfamiliar terrain of leveraging this approach between non-model species, we sought to propose hypotheses balancing biological plausibility and the possibility of novel interactions. On the one hand, we considered PPI predictions in the light of previous knowledge of protein and cellular function. However, since the molecular aspects of parasitic manipulation are still largely to be uncovered, we also considered that undescribed or unusual mechanisms may underlie these interactions. Often, we found that single proteins, especially from *Ophiocordyceps*, were in multiple host-parasite PPIs. Although in some cases these could be errant predictions, *Ophiocordyceps* effectors may indeed bind many host proteins across different pathways. This is a commonly observed phenomenon in microbe-plant interactions, and is possibly even the norm ^2, 4^. Moreover, we found that only the S8 peptidase annotation term was enriched among *Ophiocordyceps* secreted, upregulated proteins that were in PPIs with *Camponotus* proteins. This finding indicates that the PPI proteins are not a sharply distinct subset from the fungal secretome (i.e., background proteins of the enrichment analysis), which is functionally enriched for pathogenesis and proteolytic activity overall ^19^. In contrast, we detected many enriched annotations for the host side of the interactions.

### *Ophiocordyceps* proteolysis of core intracellular host proteins

Fungal PPI proteins were enriched for S8 peptidase domains, indicating that a significant number of these proteins had proteolytic functions with possibly diverse targets ^82^. Cuticular host proteins are a canonical target for such effectors as we discuss below. Additionally, multiple host kinesin motor proteins were in PPIs with these *Ophiocordyceps* proteases. Kinesins play a role in mitotic and intracellular transport processes – with key functions in vesicle transport towards the periphery of neurons ^83^. Defects in kinesin activity can lead to dysregulated neurotransmitter release at synapses, thereby contributing to impaired neurological functions and development ^83, 84^. From these predicted S8 peptidase-kinesin interactions, we infer that the fungus may be dysregulating core host cell processes. The effects of which possibly impair neuronal signaling and neurotransmitter release to contribute to modified behavioral phenotypes. We also predicted these peptidases to interact with multiple *Camponotus* nuclear pore proteins. Although some components of the nuclear pore undergo proteolytic post-translational modification to remove SUMOylation ^85, 86^, those peptidases are in a different family than S8 peptidases. If the predicted S8 peptidase-nucleopore proteins interactions are genuine, this could mean the parasite alters transport of molecules in and out of the host nucleus. Speculatively, this could facilitate entry of the fungal proteins that bind host transcription factors.

### *Camponotus* G protein-coupled receptors as targets to modify behavior

We detected over 30 ant GPCRs or GPCR subunits in PPIs with fungal proteins. Most GPCRs were from the rhodopsin/A family and included receptors related to biogenic monoamines, acetylcholine, and neuropeptides. These receptors offer a wide range of hypothetical connections to manipulated ant behavior. Commonly, these receptors and their ligands have been linked to locomotion, feeding, and circadian rhythms – processes that have been previously highlighted in hypotheses of *Ophiocordyceps* and other fungus-insect summit-disease behavioral manipulations ^10, 14, 19, 87, 88^.

The monoamine neurotransmitters dopamine, serotonin, octopamine (analogous to vertebrate norepinephrine), and tyramine have been implicated in modulating locomotor, foraging, learning, social, reproductive, and aggressive behaviors in many insects, with experimental evidence in social insects, including ants ^89–94^. Compared to the aspecific fungi tested, both a *C. floridanus* dopamine and octopamine receptor were uniquely bound by *Ophiocordyceps* upregulated secreted proteins. This interaction hypothetically implicates these host receptors in specific *Ophiocordyceps* interactions such as behavioral manipulation. We additionally detected two muscarinic acetylcholine receptors associated with sensory and motor neurons in non-insect animals ^95–97^. In bees, excessive activation of acetylcholine receptors can cause changes in locomotor, navigational, foraging, and social behaviors ^98^. Changes in such behaviors have also been observed in *C. floridanus* ants infected with *O. camponoti-floridani*. Locomotor behavior is enhanced in these ants, while social behaviors are diminished, daily timed foraging activities become arrhythmic and navigation capabilities appear less effective ^17^.

The neuropeptide GPCRs in PPIs had a range of putative ligands linking parasite interference with behavior-regulating processes such as circadian rhythm, nutritional signaling, insect hormones, and odor reception. These GPCRs included a host trissin receptor and an allatotropin receptor (annotated as an “orexin” receptor, but likely an allatotropin receptor in insects), which, compared to the aspecific fungi, only *Ophiocordyceps* was predicted to bind. Between these two receptors, dysregulation of feeding and circadian control of locomotor activity, juvenile hormone production, immune activation, and muscle stimulation are possible effects ^99–105^. Circadian rhythm is related to ant foraging behavior (and, perhaps indirectly, feeding) and is affected by *Ophiocordyceps* infection ^17^. The synchronized time-of-day summiting and biting behaviors further suggests a role for circadian control of activity ^15, 19, 23^. Interlaced with feeding and foraging processes in ants, juvenile hormone mediates insulin/feeding related signaling and social caste identity (e.g., foraging workers) ^28, 106–108^.

Among the host neuropeptide GPCR targets shared with the aspecific fungi, we predicted binding of receptors also associated with locomotion and activity levels, feeding, aggression, muscle control, juvenile hormone, reproductive behavior, or pheromones that included those activated by: allatostatin-A ^109–112^, adipokinetic hormone or corazonin (i.e., a “gondaotropin-releasing hormone” receptor) ^113^, cholecystokinin-like peptides ^114^, CCHamide-2 ^115^, and pyrokinin, pheromone biosynthesis activating neuropeptide (PBAN), and/or capa type neuropeptides ^116–118^. Odor and pheromone detection are key elements of ant communication, behavioral regulation, and social interactions ^119–121^. GPCRs typically involved in odorant response may become dysregulated by fungal effectors to dampen nestmate interaction and/or induce aberrant behavior without normal cues. Taken together, we found multiple PPIs involving ant GPCRs putatively involved in signaling systems that could control which and when certain behaviors are performed by manipulated ants.

We also predicted PPIs involving a putative host cAMP-dependent protein kinase catalytic subunit, found only between *Ophiocordyceps* and *Camponotus*, and a host G-protein subunit alpha, both of which serve important roles as components of GPCRs. This alpha subunit was also found previously to be downregulated in manipulated *Camponotus* ^19^. *Ophiocordyceps* interference with host GPCR signaling has been previously hypothesized to be mediated, at least in part, by enterotoxins and other fungal protein toxins that interfere with cAMP levels and signal transduction ^14, 19, 24, 25^.

Given the signal for GPCR interference, with both our broadest and narrowest analyses, and that many of these receptors/ligands overlap in their behavioral associations, we suggest that this provides evidence for host-parasite GPCR PPIs as one mechanism of behavioral manipulation used by *Ophiocordyceps*. The fungal counterparts in these host GPCR-parasite effector PPIs were often predicted in many interactions. These fungal proteins included five uSSPs upregulated during manipulation and correlated to manipulation in WGCNA modules ^19^. Given that these fungal proteins lack annotations, we cannot yet determine if they might be agonists or antagonists, nor whether they bind the ligand-site or elsewhere on the receptor. As these uSSPs were not predicted to bind only these host proteins, their interaction with the receptors may be less likely mediated by specific receptor-ligand binding sites than more generic contact sites elsewhere on the host protein.

### Host-pathogen interactions involving oxidation-reduction

We predicted over 300 PPIs that contributed to an enrichment of oxidation-reduction processes among the host proteins. Oxidation-reduction processes can be involved in a range of metabolic, developmental, and host-parasite interaction pathways that have been previously hypothesized to play a role in *Ophiocordyceps* infections ^14, 19, 122, 123^. Approximately half of the host proteins were putative cytochrome P450s, with enrichment signals for cytochrome P450s maintained after aspecific fungi filtering. In insects, cytochrome P450 proteins are often involved in hormone metabolism (e.g. ecdysone and juvenile hormone), stress response, xenobiotic detoxification processes, and cuticular development ^124–130^. For example, we found multiple cytochrome p450 proteins bound uniquely by *Ophiocordyceps* of the 4C1 subtype that has been implicated in stress response (e.g., social isolation, temperature, UV, or starvation) and insecticide/xenobiotic detoxification, and is regulated by hormones including juvenile hormone ^126, 131–134^. In turn, D-SCRIPT predicted that many of those host cytochrome P450s interacted with fungal cytochrome P450s. Although cytochrome P450 proteins often act independently, homo-and hetero-dimer formation of cytochrome P450 proteins can modulate protein function in positive, negative, and substrate-specific manners ^135^. Oxidation-reduction related PPIs might most strongly represent the physiological stress and antagonistic cellular processes between host and parasite. However, fungal interference with these pathways may also be critical elements of creating a host susceptible to manipulation.

### Alteration of *Camponotus* gene regulation

We found signs that suggest *Ophiocordyceps* could interfere with *Camponotus* gene regulation via proteins that bind host transcription factors and DNA-or mRNA-binding proteins. Using the host’s own cells to dysregulate behavioral pathways could be metabolically favorable for the parasite. Some of the fungal proteins in these PPIs were uSSPs and we, therefore, cannot predict the functional result of those PPIs. However, several parasite proteins were hydrolases. We suggest that these hydrolases most plausibly cleave host proteins or their post-translational modifications. Depending on the role of the protein and specific bond broken, parasite hydrolase activity could increase or decrease transcription of host genes. Among the ca. 200 host proteins in the nucleotide-binding category, we found many tied to behavioral activity via changes in locomotion, feeding/foraging, gravitaxis, light perception, circadian clocks, development and social caste related hormones, and neuronal maintenance. Covering a similar breadth of processes as the host GPCRs in PPIs, transcriptional interference may be a complementary parasite strategy to modify their hosts. However, very few of these putative targets passed the strictest shared aspecific host protein filter. As such, specific hypotheses about individual transcription factors are difficult to parse. However, as a broader functional category, many of these host proteins intersect with processes that appear relevant to manipulated ant behavior.

### Destruction of *Camponotus* structural proteins

During manipulation, the fungus also secretes proteins predicted to bind host cuticle and connective tissue proteins. Being an endoparasite, *Ophiocordyceps* has direct access to internal host connective tissues, the endocuticle layer (once the epidermis is breached), and the cuticular lining of the gut and trachea, which include tissues that contain chitin ^70, 136^. However, the fungal cell wall also contains chitin. As such, we cannot fully discount that some of these secreted fungal proteins may simply be cell wall modifying proteins that have falsely been predicted to bind chitin-associated ant proteins. Nevertheless, the fungus could be degrading cuticular proteins for nutrition, detaching muscles, or in preparation for emerging hyphae to later grow out of the host cadaver. Corroborating evidence for the disruption of the musculature stems from detailed microscopy work on *Ophiocordyceps*-infected ants at the time of manipulated summiting and biting behavior. These reports describe visual signs of fungal invasion, tissue degradation, and atrophy of manipulated ant mandible muscles ^15, 22, 29, 137^.

The predicted fungal interactors with host structural proteins included multiple proteases such as a M43-like metalloprotease and S8 subtilisin-like serine peptidase. S8 peptidases can degrade insect cuticle and are found in many fungi ^138, 139^. We also detected unannotated secreted proteins in cuticular protein PPIs that could have similar or different functions. The aspecific fungi included a generalist entomopathogen capable of degrading a wide variety of insect tissue (*C. bassiana*); consistent with this, we found no *Ophiocordyceps*-specific cuticular-protein host targets.

### Gene networks and expression broadly implicate PPIs in manipulation

*Ophiocordyceps* PPI proteins were enriched for WGCNA modules transcriptionally associated with manipulation and containing uSSPs and other previously hypothesized effectors (F1 and F2)^19^. In *Camponotus*, gene modules associated with neuronal processes and GPCRs (A14) and signaling and transcription (A10) further support the involvement of these processes with manipulation ^19^. Assuming “guilt-by-association,” in some cases this could indicate these PPIs are related to infection and manipulation processes even if we yet lack clear mechanistic hypotheses of how (e.g., PPIs with uSSPs) ^140–142^. Additionally, both upregulated and downregulated host DEGs ^19^ were enriched in predicted PPIs, possibly indicating homeostatic responses to fungal modulation of functional protein levels. As such, the proteome-level predictions that we made in this study are well in line with previous transcriptomics work. Therefore, this work serves as an additional line of evidence for certain host processes and narrows the pool of well-supported effector candidates involved with behavioral manipulation.

## Conclusion

Faced with the challenge of uncovering how a parasitism with two non-model species may operate at the molecular level, we combined multiple datatypes and tools to predict host-parasite PPIs. These approaches are feasible in many study systems, even for those that may lack the tools for sophisticated functional testing. We employed a structurally aware machine learning program, D-SCRIPT, to predict PPIs beyond well-documented interactions. By combining genomic and transcriptomic data we selected PPIs of interest with multiple bioinformatic tools to: (*i*) annotate secreted fungal proteins, (*ii*) filter interactions predicted across different species, and (*iii*) select proteins linked to increased gene expression during infection. Using the resulting PPIs, we performed enrichment analyses to identify functional categories that should be robust to computational error rates at the individual PPI level. Taken together, we produced multiple hypotheses connecting host-parasite interspecific PPIs to processes related to infection and manipulation of *Camponotus* by *Ophiocordyceps*. We highlighted evidence for PPIs involving fungal S8 proteases and PPIs involving host oxidation-reduction processes, cuticular proteins, gene regulation, and GPCRs. Especially, with analytically robust results and direct biological links to behavior, we emphasize parasitic dysregulation of GPCR signaling as a mechanism of parasitic behavioral manipulation.

## Acknowledgements

We would like to thank Biplabendu Das, who was involved in discussions of project goals and approaches, and Anna Savage, who provided constructive comments on a draft of this manuscript.

## Data Availability

The genome assemblies used for this study are available on NCBI: *O. camponoti-floridani* (GCA_012980515.1), *C. floridanus* (GCA_003227725.1), *C. bassiana* (GCA_000280675.1), *T. reesei* (GCA_000167675.2), and *S. cerevisiae* (GCF_000146045.2)^7, 19, 49, 66, 67^. Transcriptomic sequence data for *O. camponoti-floridani* and *C. floridanus* are also hosted on NCBI (BioProject PRJNA600972) ^19^. Analyzed transcriptomic data used here are available on figshare, as supplemental information associated with their original publication (https://doi.org/10.25387/g3.12121659) ^19^. Prediction and enrichment results are available in Supplementary Files S1 and S2 published with this article.

## Author Contributions

I.W., W.C.B., and C.dB. conceived the project and wrote the manuscript. W.C.B. performed the secretome annotation. I.W. performed PPI predictions, enrichment analyses, and visualization. This work was supported by the National Science Foundation (CAREER IOS-1941546 to C.dB).

## Supplementary Information

**Discussion S1:** A Word document briefly describing preliminary analysis of transmembrane fungal proteins.

**Figure S1:** Filtering aspecific PPIs and host proteins shared between *Ophiocordyceps* and alternative fungi.

**Figure S2:** D-SCRIPT edge values for tested protein pairings between extracellular fungal proteins and the ant proteome.

**Table S1:** Aspecific fungi and *Ophiocordyceps* PPI overview.

**File S1:** *Ophiocordyceps*-*Camponotus* PPI protein and enrichment data in a single Excel file with multiple tabs.

**File S2:** Aspecific fungi PPI data in a single Excel file with multiple tabs.

