## Supplemental Information for "Using machine learning to predict protein-protein interactions between a zombie ant fungus and its carpenter ant host"

**Supplementary Discussion S1**

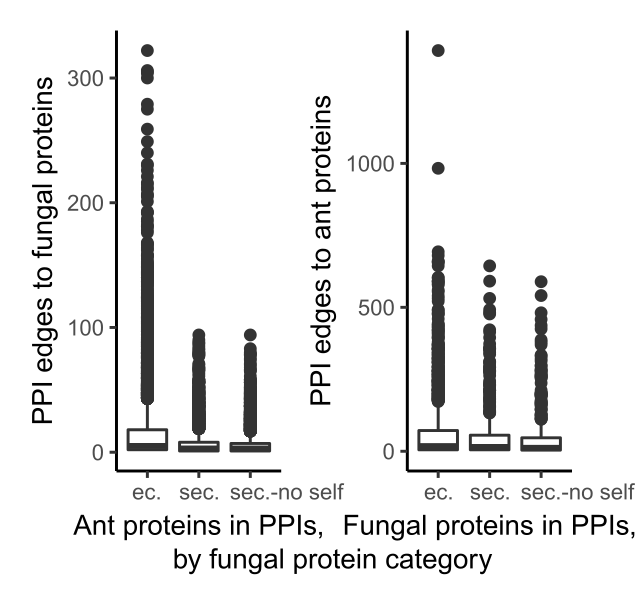
We found that *Ophiocordyceps* transmembrane proteins in these predictions to be more promiscuous binding partners compared to secreted proteins involved in possibly less informative predicted PPIs. For example, the *Ophiocordyceps* extracellular proteins with the two highest number of predicted interactions with host proteins were a transmembrane Sec61 subunit (n = 1392 PPIs) and maltose permease (n = 983 PPIs). Both proteins appear to have core cellular functions suggesting their PPI predictions may be spurious and related to their conserved roles and binding patterns. The Sec61 protein transporter plays a role in translocating proteins to the endoplasmic reticulum and the maltose permease is involved in sugar transport (58, 59). Overall, secreted *Ophiocordyceps* proteins had fewer predicted interactions (range = 1 to 644, median = 15) than transmembrane *Ophiocordyceps* proteins (range = 1 to 1392, median = 21) (Supplementary Discussion S1 Fig. 1). Similarly, the paired binding partners in *Camponotus* had somewhat fewer interactions with secreted fungal proteins (range = 1 to 94, median = 3) compared to fungal transmembrane proteins (range = 1 to 252, median = 4).

**Supplementary Discussion S1 – Figure 1. PPI connectivity after filtering predictions based on *Ophiocordyceps* extracellular category.** Box plots of the predicted number of PPIs per protein, organized by fungal protein category. PPI networks with extracellular fungal proteins (“ec.”), which include both secreted and transmembrane fungal proteins, had the highest connectivity (i.e., protein nodes with the most PPI edges). Removing PPIs involving fungal transmembrane proteins left only secreted proteins (“sec.”) and further filtering out homologous PPIs across species (“sec.-no self”) reduced connectivity. Note the difference in scale between ant protein connectivity (maximum > 300) and fungal protein connectivity (> 1,000).

**Supplementary Figure S1**

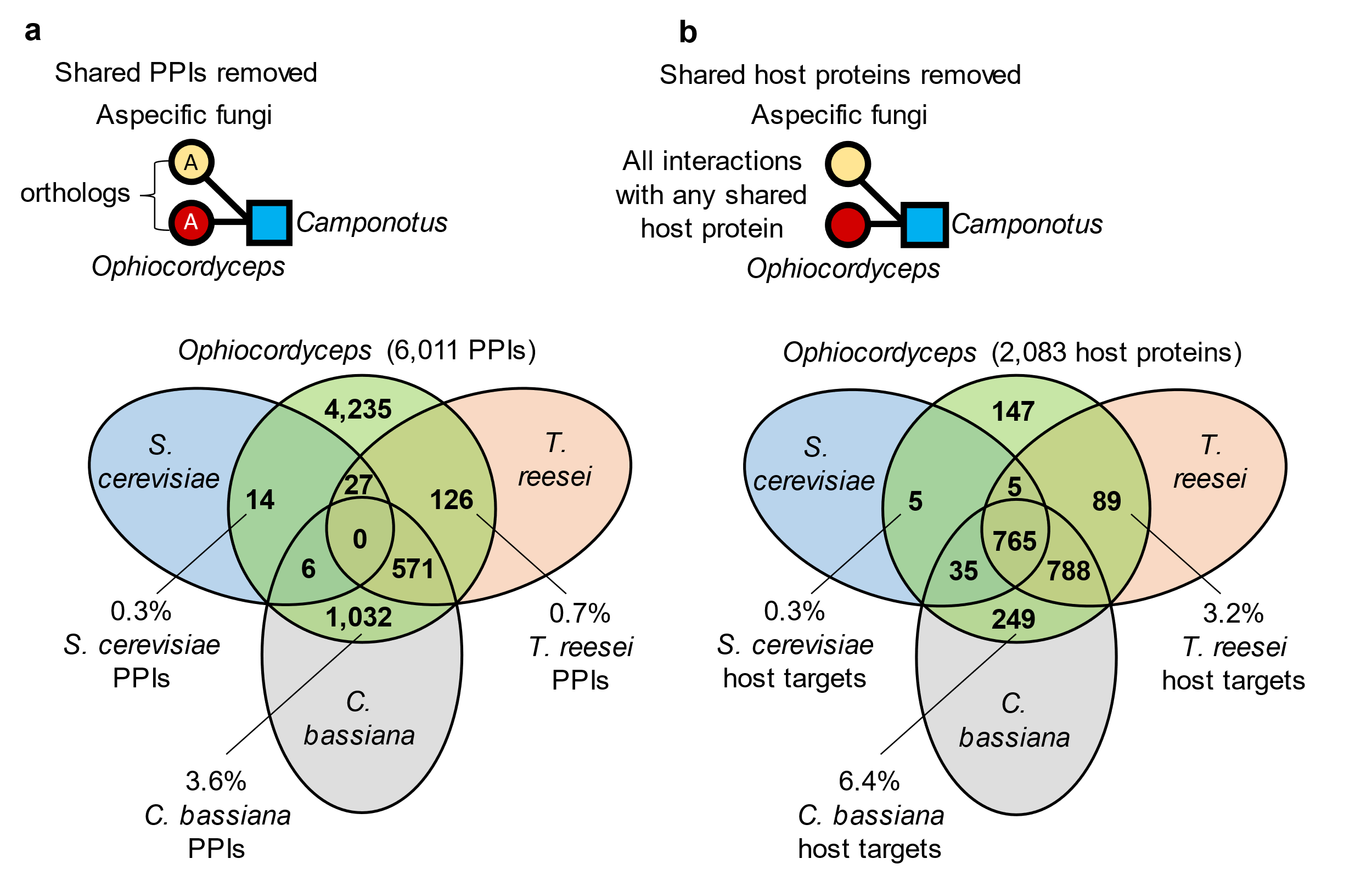
 **Supplementary Figure S1. Filtering aspecific PPIs and host proteins shared between *Ophiocordyceps* and alternative fungi.** Aspecific filtering steps were performed after selecting PPIs based on *Ophiocordyceps* secretion signals, upregulated DEGs, and no homology to self-PPIs. Both *C.* *bassiana* and *T. reesei* were similarly phylogenetically distant from *Ophiocordyceps*, with *S. cerevisiae* most distantly related. *C. bassiana* was most similar to *Ophiocordyceps* by lifestyle, being a generalist entomopathogen. a) All PPIs shared between *Ophiocordyceps* and at least one of *C. bassiana*, *T. reesei*, or *S. cervisiae* were removed. We defined aspecific homologous PPIs as those that involved a single host protein predicted to bind fungal proteins in *Ophiocordyceps* and an alternative fungus that were orthologous to each other (i.e., proteins “A” in the figure). Most PPIs were specific (4,235 of 6,011) to *Ophiocordyceps*. With most aspecific PPIs shared with *C. bassiana*. b) Removing aspecific host proteins (and therefore all PPIs and *Ophiocordyceps* proteins only predicted with these host proteins) was the strictest filter. Only 147 of the initially predicted 2,083 host proteins passed this filter. Again, most were shared with *C. bassiana*.

**Supplementary Table S1**

**Supplementary Table S1. Aspecific fungi and *Ophiocordyceps* PPI overview.** Results do not reflect any DEG or self-homology filtering steps, predictions are based on fungal secretome by host proteome. “Predicted PPIs” reflect the percent of positive predictions from D-SCRIPT relative to the total number of protein combinations tested, with the count of positive PPIs in parentheses. “Fungal proteins” is the percent of fungal proteins that were in at least one positive PPI, relative to the secretome size. “*Camponotus* proteins” is the precent of host proteins that were in at least one positive PPI, relative to the proteome size. “Fungal connectivity” shows the range of how many host proteins each fungal protein was predicted to bind and the median in parentheses. “*Camponotus* connectivity” shows the range of how many fungal proteins each host protein was predicted to bind and the median in parentheses.

| **Fungus** | **Predicted PPIs** | **Fungal proteins** | ***Camponotus* proteins** | **Fungal connectivity** | ***Camponotus* connectivity** |
| --- | --- | --- | --- | --- | --- |
| *Ophiocordyceps* | 0.33% (23,629) | 69% (402) | 27% (3,289) | 1 – 644 (15) | 1 – 94 (3) |
| *C. bassiana* | 0.28% (28,316) | 68% (567) | 23% (3,904) | 1 – 1576 (13) | 1 – 190 (3) |
| *T. reesei* | 0.27% (16,835) | 72% (365) | 23% (2,790) | 1 – 556 (12) | 1 – 131 (2) |
| *S. cerevisiae* | 0.26% (4,867) | 76% (117) | 13% (1,615) | 1 – 541 (21) | 1 – 44 (1) |

**Supplementary Figure S2**

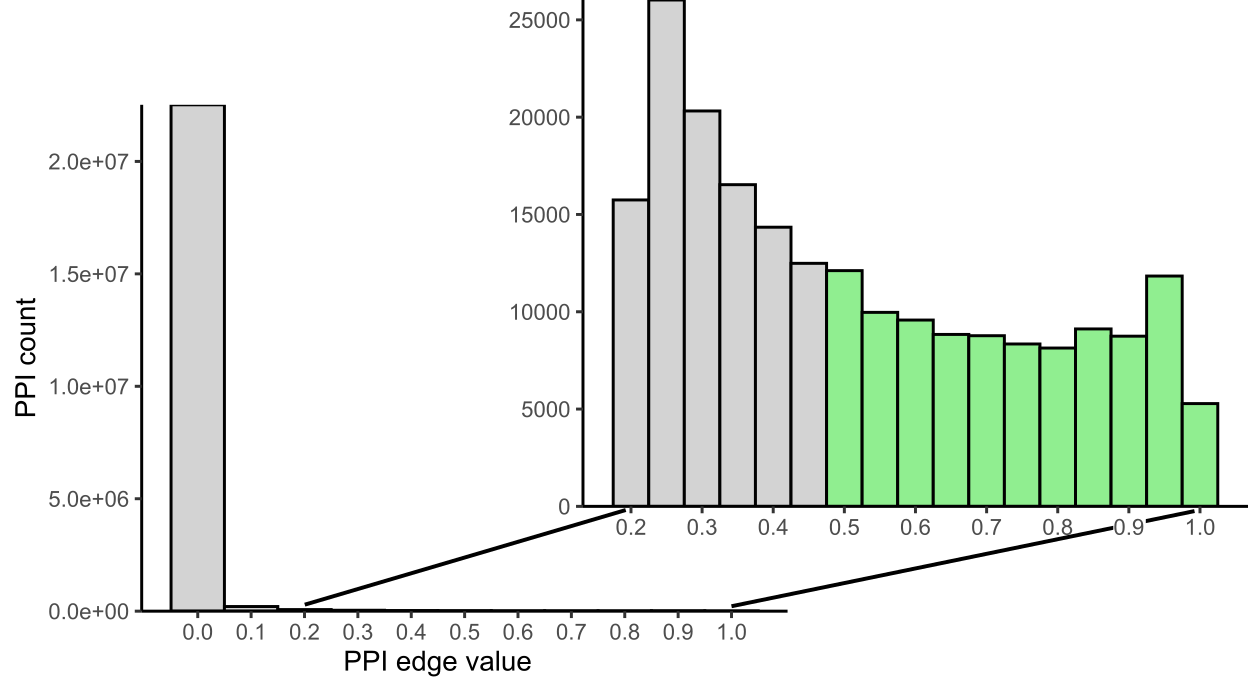

**Supplementary Figure S2. D-SCRIPT edge values for tested protein pairings between extracellular fungal proteins and the ant proteome.** D-SCRIPT edge values indicate the confidence of the model prediction for a given PPI to be true. We used the default edge value cutoff of ≥ 0.5 (green). These positive predictions comprised 0.41% of the ca. 23 million protein combinations tested. D-SCRIPT assigned most negative predictions with decisively low values far below the 0.5 threshold. As described in Supplementary Discussion S1, extracellular here refers to both secreted and transmembrane proteins.
